# Endogenous Oxytocin, Vasopressin and Aggression in Domestic Dogs

**DOI:** 10.1101/151514

**Authors:** Evan L. MacLean, Laurence R. Gesquiere, Margaret E. Gruen, Barbara L. Sherman, W. Lance Martin, C. Sue Carter

## Abstract

Aggressive behavior in dogs poses public health and animal welfare concerns, however the biological mechanisms regulating dog aggression are not well understood. We investigated the relationships between endogenous plasma oxytocin (OT) and vasopressin (AVP) – neuropeptides that have been linked to affiliative and aggressive behavior in other mammalian species – and aggression in domestic dogs. We first validated enzyme-linked immunosorbent assays (ELISAs) for the measurement of free (unbound) and total (free + bound) OT and AVP in dog plasma. In Experiment 1 we evaluated behavioral and neuroendocrine differences between a population of pet dogs with a history of chronic aggression toward conspecifics and a matched control group. Dogs with a history of aggression exhibited more aggressive behavior during simulated encounters with conspecifics, and had lower free, but higher total plasma AVP than matched controls, but there were no group differences for OT. In Experiment 2 we compared OT and AVP concentrations between pet dogs and a population of assistance dogs that have been bred for affiliative and non-aggressive temperaments, and investigated neuroendocrine predictors of individual differences in social behavior within the assistance dog population. Compared to pet dogs, assistance dogs had higher free and total OT, but there were no differences in either measure for AVP. Within the assistance dog population, dogs who behaved more aggressively toward a threatening stranger had higher total AVP than dogs who did not. Collectively these data suggest that endogenous OT and AVP may play critical roles in shaping dog social behavior, including aspects of both affiliation and aggression.

---

Aggressive behavior in dogs is a serious concern for reasons related to both public health and animal welfare. In the United States it is estimated that dogs bite ~4.5 million Americans annually, with approximately half of these bites directed toward children (Centers for Disease Control and Prevention, 2003; Gilchrist, Sacks, White, & Kresnow, 2008). In addition to this impact on human lives, aggression (toward humans or other dogs) is also one of the most common reasons that dogs are relinquished to animal shelters (Salman et al., 1998), with ~2 million of these dogs being euthanized every year (Patronek, Glickman, Beck, McCabe, & Ecker, 1996). Despite these widely recognized concerns, we know relatively little about the biological factors underlying dog aggression.

Previous studies on the biology of canine aggression have focused predominantly on the role of androgens and the serotonergic system. Although testosterone is positively associated with aggression in many species (Archer, 1988), studies of androgens and aggression in dogs have been largely inconclusive. For example, in some studies gonadectomy (which yields decreased androgen production) has been linked to a mild reduction in male dog aggression toward both other dogs and people (Neilson, Eckstein, & Hart, 1997), whereas in others neutered dogs were found to be more aggressive (Guy et al., 2001). Findings on the serotonergic system have been more consistent than those for androgens. Specifically, some dogs with a history of aggression are characterized by low levels of serotonin or serotonin metabolites – in both blood and cerebrospinal fluid (CSF) – and this finding is especially pronounced in lineages prone to aggression (Amat et al., 2013; Haug, 2008; León et al., 2012; Reisner, Mann, Stanley, Huang, & Houpt, 1996; Rosado et al., 2010). Because of the inhibitory effect of serotonin on aggression, one common intervention for aggressive dogs has been to increase serotonin availability through selective serotonin reuptake inhibitors (SSRIs; Haug, 2008). Although testosterone and serotonin may both have important roles in regulating aggression, research with other mammalian species indicates that oxytocin and arginine vasopressin also play major roles in the inhibition and facilitation of aggressive behaviors (Albers, 2012; Caldwell, Lee, Macbeth, & Young III, 2008; Carter, 1998). However, few studies have investigated the links between these neuropeptides and aggressive behavior in dogs.

Oxytocin (OT) and arginine vasopressin (AVP) are closely related nonapeptides with wide ranging effects on social behavior, cognition, and stress responses (Carter, 1998; Carter, Grippo, Pournajafi-Nazarloo, Ruscio, & Porges, 2008; Donaldson & Young, 2008; Goodson & Bass, 2001). Although the biological effects of OT and AVP can be similar in many cases, in others they are antagonistic (Neumann & Landgraf, 2012). With respect to affective states and social behavior, OT inhibits the sympathoadrenal axis, reduces anxiety, and can promote affiliative behavior. In contrast, AVP increases sympathoadrenal activity, is anxiogenic, and in some cases facilitates aggression (Carter, 1998; Ferris, 1992). However, both peptides can have effects that are sex- and species-specific, and depend on site of action in the brain, as well as characteristics of the receptor (Kelly & Goodson, 2014). Moreover, both peptides are capable of binding to one another’s receptors, and the dynamic balance between OT and AVP is hypothesized to mediate a wide spectrum of emotional states and social behaviors (Neumann & Landgraf, 2012).

Recent studies with dogs have highlighted the role of OT in affiliative behavior and positive affective states. For example, dogs exhibit an increase in OT after friendly interaction with a human (Nagasawa et al., 2015; Odendaal & Meintjes, 2003; Rehn, Handlin, Uvnäs-Moberg, & Keeling, 2014), or other pleasurable experiences (Beetz, Uvnäs-Moberg, Julius, & Kotrschal, 2012; Mitsui et al., 2011). Recently, polymorphisms in the oxytocin receptor gene (*OXTR*) have been linked to human-directed social behavior in dogs (Kis et al., 2014; Oliva, Wong, Rault, Appleton, & Lill, 2016), and dogs treated with intranasal OT have been documented to exhibit increased affiliative behavior toward both humans and other dogs (Nagasawa et al., 2015; Romero, Nagasawa, Mogi, Hasegawa, & Kikusui, 2014; but see Hernádi et al. 2015). Lastly, OT administration has been documented to enhance some aspects of dog-human communication (Oliva, Rault, Appleton, & Lill, 2015), including cognitive skills that may be convergent between humans and dogs (MacLean & Hare, 2015; MacLean, Herrmann, Suchindran, & Hare, 2017). Thus, current data suggest that OT both facilitates and responds to some types of affiliative and cooperative social interaction in dogs.

Although no studies have investigated the role of AVP in dog aggression, data from other mammalian species suggest that AVP plays an important role in regulating aggression toward unfamiliar individuals. For example, early studies on AVP and aggression revealed that microinjection of AVP into the hypothalamus of golden hamsters led to increased aggression toward unfamiliar conspecifics, whereas hamsters receiving an AVP antagonist displayed a dose-dependent decrease in biting and latency to attack unfamiliar individuals (Albers, 2012; Ferris, 1992; Ferris et al., 2006; Ferris et al., 1997; Ferris & Potegal, 1988). Although these findings have been replicated in several other species (Bester - Meredith, Martin, & Marler, 2005; Gobrogge, Liu, Jia, & Wang, 2007), other experiments reveal that AVP can both facilitate or inhibit aggression, depending on the site of action in the brain or sex-specific factors (reviewed in Albers, 2015; Kelly & Goodson, 2014), and AVP may be critical for some forms of affiliative behavior (Carter, Devries, & Getz, 1995). In contrast to these rodent studies which have addressed localized functions of AVP, human studies have measured AVP in cerebrospinal fluid (CSF) or the periphery to assess potential links between overall circulating levels of AVP and social behavior. With respect to aggression, Coccaro et al. (1998) measured AVP in human CSF and found positive associations between AVP and a life history of aggression, and studies administering intranasal AVP in men led to decreased perceptions of friendliness in unfamiliar faces (Thompson, George, Walton, Orr, & Benson, 2006).

Taken together, these findings suggest that OT may play a larger role in affiliative social behavior, anxiolysis, and the inhibition of aggression, whereas AVP – though also critical to bond formation and parental behavior – may play a larger role in anxiogenesis and aggression. To investigate the links between OT, AVP and aggressive behavior in dogs we conducted two studies in which dogs were individually exposed to various stimuli: (1) three-dimensional dog models, (2) video images of other dogs, and/or (3) a threatening human, and we recorded the resulting aggressive responses. Free (unbound) and total (free + bound) plasma OT and AVP concentrations were also determined and used as predictors of behavior in these contexts. We first conducted a series of methodological studies to validate sample preparation protocols for the measurement of OT and AVP in dog plasma. In Experiment 1, we compared the behavior and OT/AVP concentrations of two group of dogs: a ‘case group’ - dogs recruited because of their known history of aggression toward unfamiliar conspecifics - and a ‘control group’ - dogs with no previous history of aggression toward conspecifics, who were matched to cases on the basis of breed, sex, and age. These dogs were exposed to life-like three-dimensional dog models, as well as video-projected stimuli featuring dogs engaged in a variety of non-aggressive behaviors. In Experiment 2 we compared the hormone concentrations of a population of assistance dogs – who have been selectively bred for affiliative and non-aggressive behavior – and the companion dogs tested in Experiment 1. We also tested the assistance dogs with the video stimuli used in Experiment 1, as well as in a temperament evaluation during which dogs were exposed to a life-like three-dimensional dog model, and an unfamiliar human who approached the dog in a threatening manner.

## OT and AVP method validation

Although immunoassays for OT and AVP have been available for decades, there are ongoing debates regarding whether sample extraction is needed previous to the assay for accurate peptide quantification (Christensen, Shiyanov, Estepp, & Schlager, 2014; Leng & Sabatier, 2016; e.g., Robinson, Hazon, Lonergan, & Pomeroy, 2014; Szeto et al., 2011). Sample extraction is typically performed to isolate the analyte of interest and to limit cross-reactivity during the assay. However, extraction procedures also have the potential to remove biologically relevant products leading to erroneously low estimates of the target analyte (Carter, 2014; Martin & Carter, 2013; Martin, 2014). Because these methodological considerations have not been systematically investigated in dogs, we first conducted a series of studies to determine the most appropriate methods for the analysis of peptide hormones in dog plasma. Details of these studies are reported in the supplemental information (SOM) and the results are briefly summarized below. Because recent studies demonstrate that OT (and likely AVP) readily bind to proteins in plasma, we explored techniques measuring both free (unbound), and total (free + bound) peptide concentrations.

## Method and Results

Oxytocin samples were assayed using commercially available enzyme-linked immunosorbent assay (ELISA) kits from Arbor Assays (K048) and Cayman Chemical (500440). The Arbor Assays kit was used for all analyses with the exception of the measurement of free OT with the assistance dog population in Experiment 2. This change was implemented because free OT concentrations measured with the Arbor Assays kit in Experiment 1 were near the lower limit of detection, and subsequent analyses in our lab revealed that free OT in dog plasma was detectable in a better region of the standard curve with the Cayman Chemical kit. All vasopressin samples were assayed using a commercially available ELISA kit from Enzo Life Sciences (ADI-900-017A).

### Measurement of Free OT and AVP

We assessed whether free OT and AVP concentrations can reliably be measured in dog plasma by assessing parallelism, accuracy and precision of the assay, using a pooled sample of dog plasma that was or was not extracted prior to the assay through solid phase extraction (SPE).

Serial dilutions of the plasma pool showed that extracted samples yielded better parallelism than non-extracted samples for both OT and AVP (see SOM). For a series of dilutions using extracted samples, linear models predicting the observed concentration as a function of the expected concentration had slopes of ~1 and intercepts that did not differ from the expected value of 0 (OT: β = 1.05, p = 0.50; AVP: β = 0.91, p = 0.76). In contrast, dilutions of non-extracted samples had slopes >1 that did not pass through the origin (OT: β = 1.55, p < .01; AVP: β = 1.59, p = 0.05). Assay accuracy – assessed by spiking kit standards with extracted and non-extracted pools – was better with extracted samples for OT (mean recoveries: extracted = 118%, non-extracted = 153%), but better with non-extracted samples for AVP (mean recoveries: extracted = 69%, non-extracted = 104%). Based on the superior parallelism and accuracy following SPE extraction, we opted to extract all samples for analyses of free peptide concentrations. However, we also analyzed samples from Experiment 1 without extraction to compare the results of both approaches (SOM). AVP concentrations for the same samples analyzed with and without extraction were not correlated (t_114_ = 1.26, p = 0.21, R = 0.12), while OT concentrations measured by the two methods exhibited a modest negative correlation (t_114_ = −2.45, p = 0.02, R = −0.22). For the extracted plasma pool the intra-assay CVs from 6 replicates were 7.7% for OT (mean concentration = 22.7 pg/mL) and 10.1% for AVP (mean = 5.3 pg/mL). Inter-assay CVs across 4 assays were 13.9% for OT (mean = 16.2 pg/mL) and 9.2% for AVP (mean = 4.4 pg/mL).

### Measurement of Total OT and AVP

Recent data suggest that OT binds strongly to plasma proteins which may prevent its detection in plasma (Brandtzaeg et al., 2016; Martin & Carter, 2013; Martin, 2014). Given its structural similarity to OT and the presence of a disulfide bridge, it is likely that AVP exhibits similar binding patterns. Indeed, most studies employing SPE followed by ELISA or radioimmunoassay for the measurement of free OT/AVP report concentrations that are near the limits of detection, and similar results have been obtained using liquid chromatography mass spectrometry (Johnsen, Leknes, Wilson, & Lundanes, 2015; Zhang, Zhang, Fast, Lin, & Steenwyk, 2011). We have recently shown that a reduction / alkylation and protein precipitation (R/A PPT) procedure – which liberates bound OT from plasma proteins – allows for the detection of much higher concentrations of OT, and have validated this approach with dog plasma analyzed by ELISA (Brandtzaeg et al., 2016). Due to the protein precipitation step, this process also eliminates the matrix interferences commonly observed when working with neat plasma. Here we used this approach for both OT and AVP (see SOM). Samples extracted via SPE should measure ‘free’ peptide concentrations, reflecting acute activity at the time of the study, whereas samples prepared via R/A PPT should represent total OT and AVP concentrations, and provide a biomarker of longer-term individual differences (free and bound concentrations; Brandtzaeg et al., 2016).

## Experiment 1

Experiment 1 was a case-control study in which dogs with a history of aggression (hereafter cases) toward unfamiliar dogs while walking on leash (“leash aggression”) were compared to a matched control group (hereafter ‘controls’) with no history of aggression. All testing took place at the Veterinary Health and Wellness Center at North Carolina State University’s College of Veterinary Medicine.

## Method

### Subjects

Cases were recruited through dog-related email list-serves and area dog trainers specializing in cases of leash aggression. Recruitment materials solicited owners with dogs who routinely snarl, growl or lunge at unfamiliar dogs while on leash. Individuals expressing an interest in the study participated in an initial phone screening to verify that their dog exhibited a chronic pattern of aggression toward other unfamiliar dogs, and met the inclusionary criteria described below. When cases met the inclusionary criteria, matched controls (based on breed, sex, and age) were recruited from a database of pet owners maintained by the Duke Canine Cognition Center. Owners of control dogs participated in an initial screening to verify that their dog was not aggressive toward unfamiliar conspecifics, and that dogs met the basic inclusionary criteria. All owners were offered a free veterinary exam, complete blood count (CBC), and serum biochemistry panel for their dog in exchange for participation. To be eligible for participation subjects were required to be between 1-9 years of age, 4.5-70 kg in weight, spayed or neutered, and to have an up-to-date rabies vaccination. Dogs were excluded from participation if they had chronic illnesses, a history of aggression toward humans or familiar dogs within the household, abnormal results for the CBC or chemistry panel, or had received psychoactive medications within the past 30 days. In total, 46 dogs began the study, but three subjects were subsequently excluded either because blood could not be collected during the veterinary exam (N = 2), or due to abnormalities in the CBC results (N = 1). Subject demographics for matched cases and controls are shown in Table 1. All dog owners signed informed-consent documents prior to participation and testing procedures adhered to regulations set forth by the Institutional Animal Care and Use Committee at North Carolina State University (Protocol#: 14-184-O).

**Table 1.** Subject demographics for Experiment 1.

| Breed | Group | Case / Control | Sex | Age (years) |
| --- | --- | --- | --- | --- |
| German Shepherd | A | case | F | 6 |
| German Shepherd | A | control | F | 9 |
| Beagle mix | B | case | F | 7 |
| Beagle | B | control | F | 4 |
| American Staffordshire Terrier mix | C | case | M | 4.5 |
| American Staffordshire Terrier mix | C | control | M | 4 |
| Weimaraner | D | case | M | 6 |
| Golden Retriever | D | control | M | 7 |
| Golden Retriever | E | case | F | 3 |
| Golden Retriever | E | case | F | 4 |
| Potcake Dog | F | case | M | 2 |
| Labradoodle | F | control | M | 4 |
| American Staffordshire Terrier | G | case | F | 2 |
| American Staffordshire Terrier mix | G | control | F | 3 |
| Lhasa Apso mix | H | case | M | 8 |
| Cavalier King Charles Spaniel | H | control | M | 7 |
| Papillon | H | control | M | 6 |
| German Shepherd | I | case | M | 4 |
| German Shepherd | I | control | M | 4 |
| Hound mix | J | case | M | 6 |
| American Staffordshire Terrier mix | J | case | M | 5 |
| Treeing Walker Coonhound | J | control | M | 6 |
| Corgi mix | K | case | F | 3 |
| Corgi mix | K | control | F | 6 |
| Australian Cattle Dog mix | L | case | F | 3 |
| Border Collie mix | L | control | F | 3 |
| Boxer | M | case | M | 5 |
| Boxer | M | control | M | 3 |
| Poodle (Standard) | N | case | M | 5 |
| Mastiff mix | N | case | M | 4.5 |
| Poodle (Standard) | N | control | M | 6 |
| Terrier mix | O | case | M | 6 |
| Australian Cattle Dog | O | case | M | 6 |
| Border Collie mix | O | control | M | 6 |
| Golden Retriever | P | case | F | 4 |
| Golden Retriever | P | control | F | 3 |
| Greyhound | Q | case | M | 4 |
| Greyhound | Q | control | M | 2.5 |
| Labrador Retriever | R | case | M | 8 |
| Labrador Retriever | R | control | M | 9 |
| Australian Cattle Dog | S | case | M | 5 |
| Australian Cattle Dog | S | control | M | 8 |
| Australian Sheparad mix | T | case | M | 7 |
| Australian Cattle Dog | T | control | M | 4 |

### Apparatus and Stimuli

A schematic of the testing room is shown in Figure 1. The test room was divided into two sections by a wall of filing cabinets (1.22m high) with a 1.7m gap in this wall to allow subjects to view stimuli presented in the other half of the room. An opaque curtain was hung across this gap so that subjects could see through to the other side of the room only during periods of stimulus presentation. Owners sat in a chair 2.5m from the dividing wall with a rubber mat adjacent to this chair where dogs were positioned at the start of the study. All dogs wore a 1.25m leash attached to a neck collar to allow them to move freely within a fixed radius from their starting position during the test. Video stimuli were projected with an NEC video projector (model VT695; only present during video trials) on a 1m x 1.5m white poster board that was positioned 1.5m behind the opening in the room divider. Audio speakers were positioned behind this poster board so that sounds were presented from directly behind the area where visual stimuli were presented. All trials were filmed with two high definition camcorders, one positioned at the back of the room, which captured the stimulus presentations and dog (from the rear), and the other from the gap in the center of the room, which captured the dog’s behavior (oriented toward the dog’s face). This second camera was the primary angle used for all behavioral coding.

**Figure 1.**
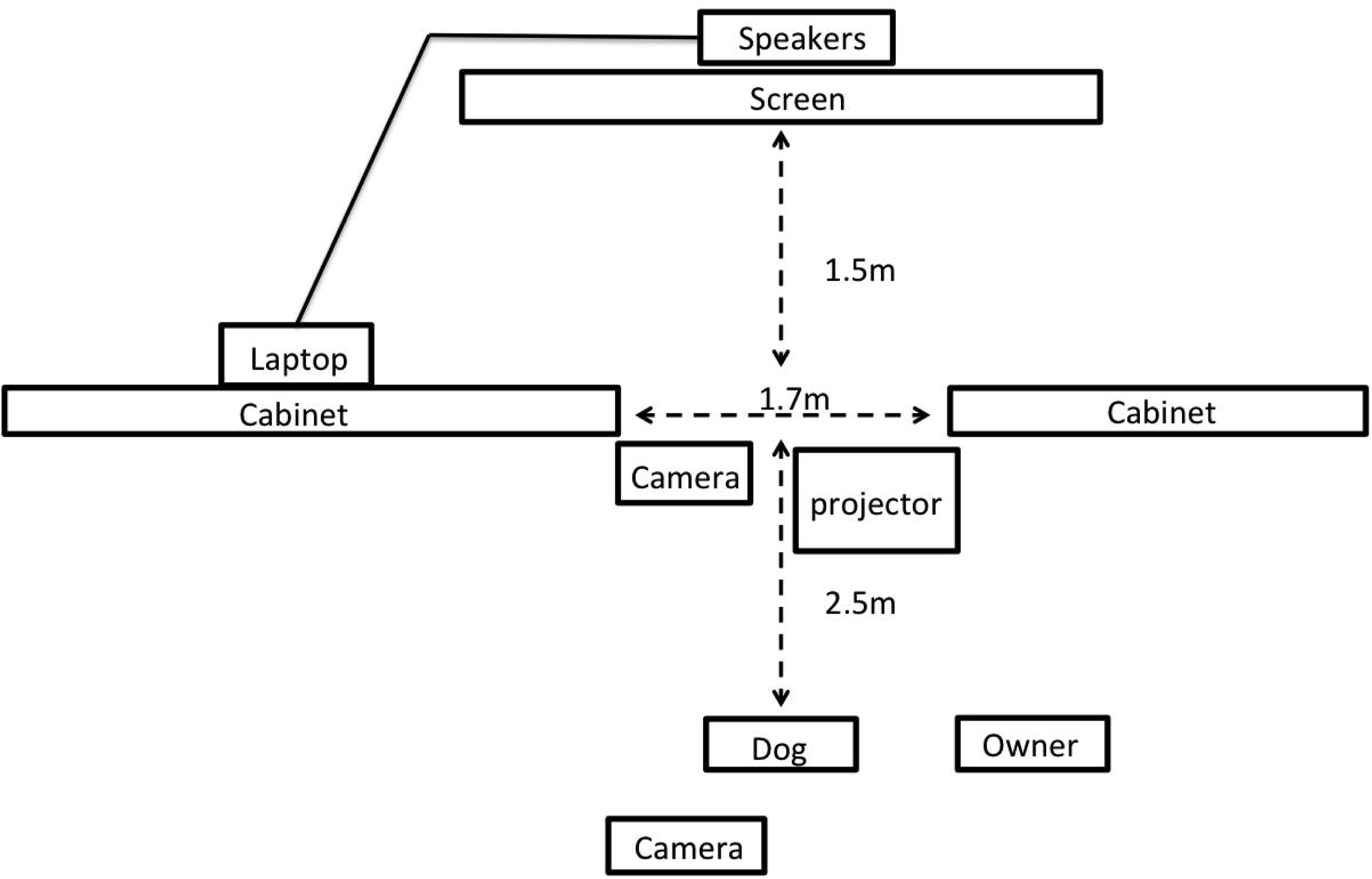

Three-dimensional stimuli consisted of life-like dog models of three different sizes (small – Jack Russell Terrier; medium – Shetland Sheepdog; large – Old English Sheepdog) and three inanimate objects (control stimuli) of comparable size (yellow box, black trash bag filled with paper, inflated blue yoga ball). Prior to the presentation of each stimulus we played a brief (~2s) auditory stimulus to attract the dog’s attention to the area in which the visual stimulus would be presented. For dog models these stimuli were barking sounds appropriate for the specific model’s body size, and for control stimuli these sounds were arbitrary sound effects. Video stimuli (each 15s long) consisted of footage from DOGTV (a company which produces digital content optimized for dog vision). Dog clips included images of a dog being walked on leash, a dog resting in the grass, and two dogs playing. Control clips included images of water running across rocks, and panoramas of a forest canopy or flowers. Audio content was added to these clips such that all control clips had soft instrumental music, and dog clips included the sound of the dog panting, or of dog play vocalizations. The audio for dog clips was digitally edited so that the sounds and visual images were synchronous.

### Procedure

#### Veterinary Exam and Background Information

After scheduling their research visit, all participants were provided with an electronic link to complete the Canine Behavioral and Research Questionnaire (C-BARQ), a validated and reliable instrument that measures various aspects of canine behavior, and has been used in previous studies of canine aggression (Duffy, Hsu, & Serpell, 2008; van den Berg, Heuven, van den Berg, Duffy, & Serpell, 2010). Upon arrival, clients were escorted to a quiet consultation room where a veterinary technician collected background information about the dog’s behavioral and medical history, and clients signed informed consent documentation. Dogs were then taken to a nearby exam room and the attending veterinarian performed a physical examination and collected the first blood sample to be used for a CBC, chemistry panel, and analysis of free and total OT and AVP. At the conclusion of the exam, dogs and their owners were brought to the test room where the primary experimenter (E1) explained the procedure and provided instructions for handling the dog during the test. Owners were asked to hold the dog’s leash against a fixed point on the arm of their chair, and to refrain from interacting with their dog during the test trials, regardless of the dog’s behavior.

#### Three-dimensional Stimuli

At the start of each trial, the curtain was closed preventing dogs from viewing activity on the experimenter’s side of the room. E1 positioned himself behind the curtain with the stimulus, and a second experimenter (E2) played the audio clip to attract the dog’s attention. E2 then opened the curtain with a pulley, and dogs observed E1 and the stimulus for 15 seconds at which point the curtain was closed. For dog stimuli, E1 held a leash that was looped around the model dog’s neck, and gently petted the dog. For control stimuli, E1 performed comparable motions (e.g. touching or patting the box and bag, or rotating and lightly bouncing the ball). E1 looked at the stimulus throughout the trial and avoided eye contact with the subject. Six trials were conducted and the stimulus type (dog model, control object) alternated between trials. We used two fixed orders for the stimulus presentation (order 2 was the reverse of order 1) and the stimulus order was counterbalanced within groups (case and control) and consistent within matched case-control pairs. Examples of the procedure and subject responses are shown in Movie S1 (click for web version).

#### Video Stimuli

At the conclusion of the final three-dimensional stimulus the curtain was left open, E1 left the testing area and started the video. Like the three-dimensional stimuli, the order of video stimuli alternated between clips featuring dogs, and control clips (nature scenes), and we used two orders of stimuli presentation that were counterbalanced as described above. Each clip was separated by 10s during which the screen was black. At the conclusion of the final video stimulus dogs were taken to a nearby exam room and a second blood sample was collected to assess changes in free OT and AVP after exposure to the test stimuli.

#### Scoring and Analysis

From video we coded each trial for the duration of barking and growling, as well as the number of times that the dog lunged at the stimulus, or raised her upper lip in an aggressive display. Raised hackles (erectile hairs along the back of the dog) could not be coded due to heterogeneity in coat type in this diverse sample, which precluded comparable measures across subjects. All trials were coded by two independent observers blind to the hypotheses and inter-rater agreement was excellent for all measures (bark: R = 0.95; growl: R = 0.98; lunge: R = 0.95, raise lip: R = 0.89). Preliminary analyses revealed much stronger reactions to the three dimensional stimuli than the video stimuli, with very few dogs exhibiting aggressive responses to the latter (Table 2). Therefore, our analyses of behavior during the test were restricted to trials incorporating the three-dimensional stimuli. Prior to analysis, each subjects’ scores were averaged across the three test trials involving dog stimuli (i.e. excluding trials with control objects) and all data were standardized by conversion to z-scores (within each behavioral category). Because a raised-lip display was observed in only one dog, this variable was dropped from analysis. For the purpose of generating a composite index of aggressive behavior, we conducted a principal components analysis with the z-scores for barking, growling and lunging. The first principal component was loaded positively by all three variables, accounted for 54% of variance, and scores for this component were used as the primary measure of aggression (hereafter ‘composite aggression score’).

**Table 2.** Behavioral differences between the test (dog) and control (non-dog) conditions with three-dimensional and video-projected stimuli. Dogs behaved more aggressively toward the conspecific models when presented with three-dimensional, but not video stimuli. Lunging was not observed in response to the video.

|  | behavior | t | p |
| --- | --- | --- | --- |
| 3D stimuli | bark | -2.42 | .02 |
|  | growl | -2.38 | .02 |
|  | lunge | -3.08 | < 0.01 |
| Video stimuli | bark | -1.03 | .31 |
|  | growl | -1.51 | .14 |
|  | lunge | - | - |

Hormonal predictors of case-control status were assessed using conditional logistic regression (Gail, Lubin, & Rubinstein, 1981). Unless otherwise noted, comparisons of behavior between the case and control groups were implemented using linear or generalized linear mixed models with the pair identifier for each matched-pair as a random effect. For hormonal analyses we included sex, age, body mass and assay plate as covariates, and for behavioral analyses we included sex, age, and body mass as covariates. In all cases, individual model predictors were evaluated in a model-comparison framework using a likelihood ratio test to determine the change in likelihood when individual variables were added to the model. Hormone data were log transformed prior to analysis.

#### Sample Collection and Hormone Analysis

All blood samples were collected into vacutainers containing K3 EDTA, centrifuged for 20 minutes at 3000 RPMs, and the separated plasma was divided into 1mL aliquots and frozen at −80 °C until assay. Samples from matched cases and controls were run on the same plate. We analyzed free OT and AVP in samples collected both before and after the experiment in order to assess short-term changes in peptide release. However, pre and post samples were highly correlated (OT: R = 0.69; AVP: R = 0.67) with no significant changes across time in either group (SOM). Therefore, we report the results only from the pre-test samples below, and additional analyses involving post-test samples, and changes across time are reported in the SOM. Total OT and AVP were measured only in pretest samples as this measure was intended to provide a longer-term and more stable measure of individual differences.

## Results

### Behavior

Overall, dogs exhibited significantly more time barking and growling, and lunged at the stimulus significantly more frequently when three-dimensional dogs were presented in comparison to three-dimensional control objects (paired t-tests, Table 2). In contrast, video stimuli provoked very few responses in either condition, and behavior did not vary depending on whether the video featured another dog, or control content (Table 2). Therefore, analyses of behavior during the experiment focus exclusively on the trials incorporating three-dimensional stimuli.

Cases and controls differed significantly in their responses to the dog stimuli with cases exhibiting more barking, growling, and lunging than controls, as well as higher composite aggression scores (Figure 2). These behavioral differences were specific to test trials (in which dog models were presented) and the behavior of the groups did not differ during control trials (bark: χ2 = 1.30, df = 1, p = 0.25; growl: χ 2 = 1.34, df = 1, p = 0.25; lunge: χ 2 = 0.04, df = 1, p = 0.84).

**Figure 2.**
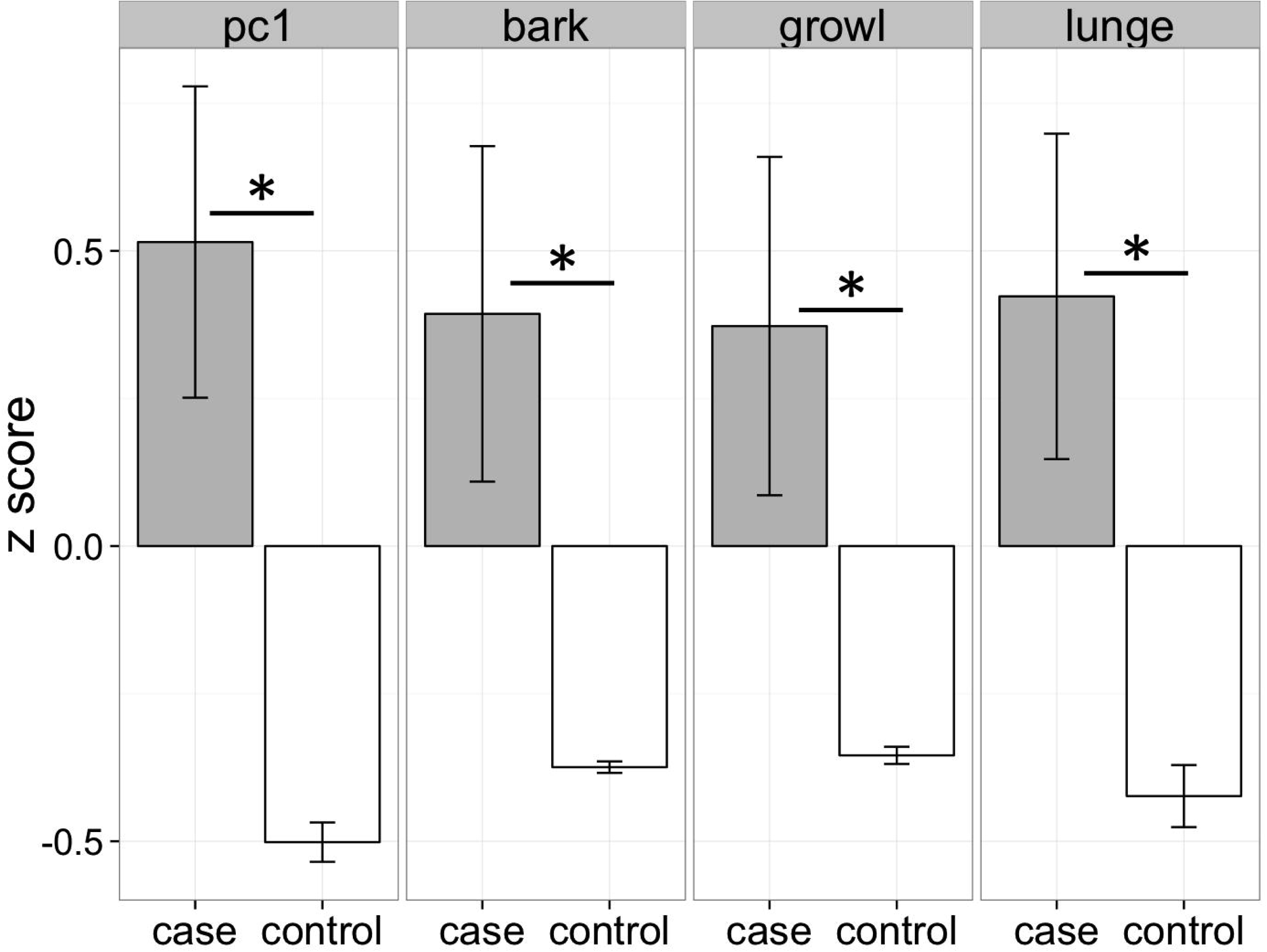

#### Free Oxytocin and Vasopressin

A conditional logistic regression revealed that cases had significantly lower free AVP levels prior to the test than controls (Figure 3; χ 2 = 10.20, df = 1, p < 0.01), but that free OT concentrations did not differ between groups (Figure 3; χ 2 = 0.01, df = 1, p = 0.94). Cases tended to have higher free OT:AVP ratios, but this difference was not significant (χ 2 = 2.86, df = 1, p = 0.09). Using the C-BARQ as a measure of aggression in the real world, we observed similar findings. Specifically, dogs reported to be more aggressive toward other dogs were characterized by significantly lower levels of free plasma AVP (SOM).

**Figure 3.**
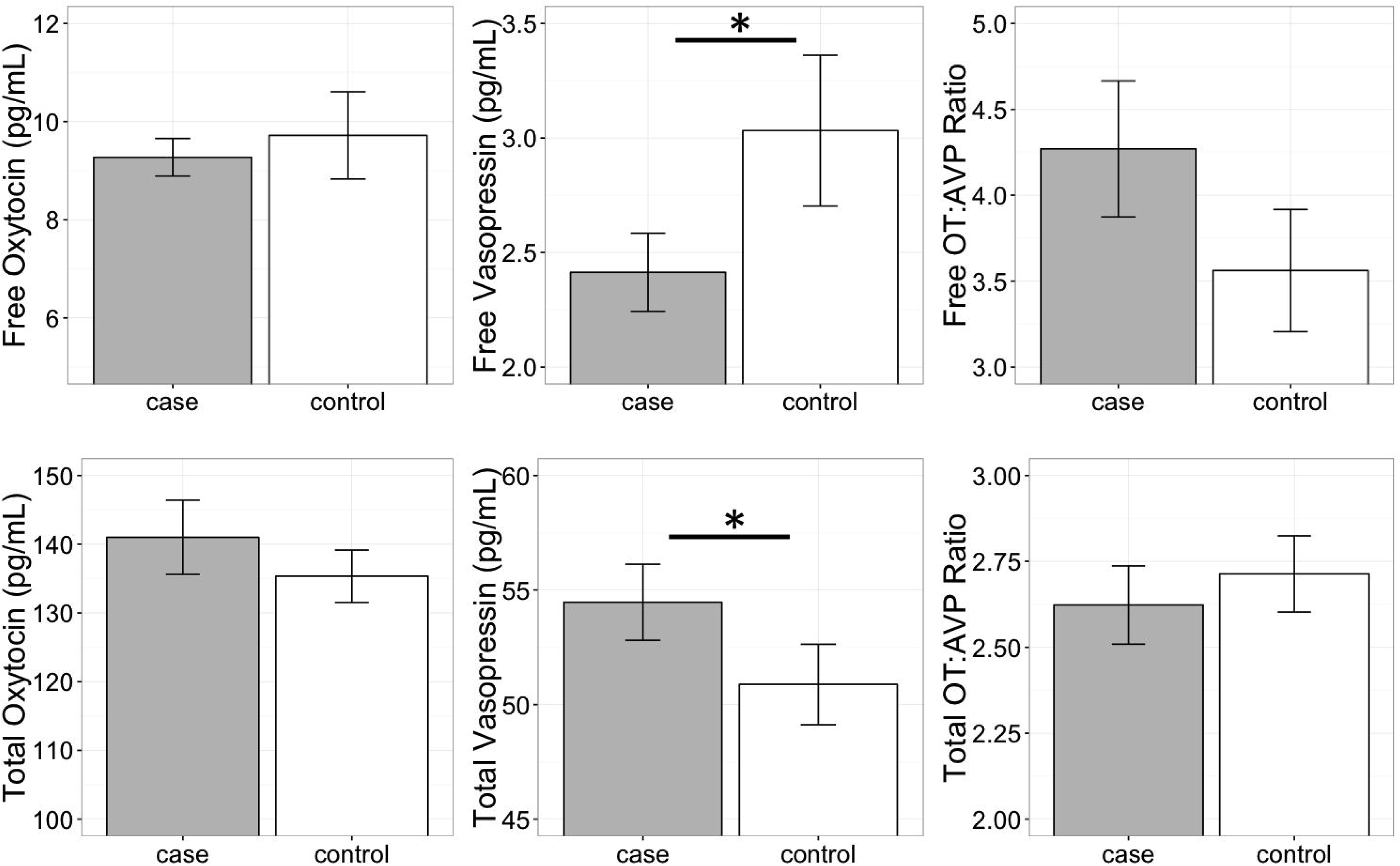

#### Total Oxytocin and Vasopressin

In contrast to the analysis of free peptide concentrations, a conditional logistic regression revealed that cases had significantly higher total AVP than controls (Figure 3; χ 2 =5.42, df = 1, p = 0.02). Neither total OT, nor the total OT:AVP ratio differed between groups (total OT: χ 2 = 0.01, df = 1, p = 0.91; total OT:AVP ratio: (χ 2 = 0.74, df = 1, p = 0.39). Lastly, using total peptide concentrations as predictors of C-BARQ scores, total AVP was positively associated with higher levels of dog-directed aggression (SOM).

## Discussion

Cases and controls differed significantly in their reactions to the three-dimensional models, with cases exhibiting higher levels of aggression than controls. Because similar behavioral differences were not observed when subjects were presented with control stimuli, this suggests that aggressive responses were both specific and social in nature, and presumably triggered by the verisimilitude of the models. However, it is noteworthy that not all cases reacted aggressively toward the dog models (SOM), despite the fact that these dogs were recruited specifically due to owner-reports of chronic aggression toward unfamiliar conspecifics. Therefore, while model dogs may provide a useful tool for research purposes, response to these inanimate models is unlikely to correlate perfectly with real-world behavior (Shabelansky, Dowling-Guyer, Quist, D’Arpino, & McCobb, 2015).

AVP (but not OT) concentrations prior to the test differed significantly between cases and controls. However, the direction of this difference depended on whether only the free portion, or total peptide concentrations were considered, an issue we revisit below. Specifically, cases were characterized by lower levels of free AVP, but higher levels of total AVP. In contrast, neither OT measure differed significantly between cases and controls. This finding lends further support to the idea that while structurally and functionally related, AVP may be more strongly associated with aggression than OT.

## Experiment 2

Experiment 2 evaluated hormonal predictors of variance in social behavior in a population of candidate assistance dogs tested at Canine Companions for Independence (CCI) in Santa Rosa, CA. Additionally, we compared endogenous OT and AVP levels between this population and the population of pet dogs studied in Experiment 1. Because this assistance dog population has been under active selection for friendly and non-aggressive temperaments for > 40 years, we expected that if OT and AVP play critical roles in shaping these traits, then this population may exhibit unique neuroendocrine characteristics relative to a population of pet dogs.

## Method

### Subjects

Thirty candidate assistance dogs from CCI participated in Experiment 2. Demographic information for all subjects is shown in Table 3.

**Table 3.** Subject Demographics for Experiment 2.

| Breed | Sex | Age | Intact |
| --- | --- | --- | --- |
| Lab - Golden Cross | M | 1.8 | no |
| Lab - Golden Cross | F | 1.6 | yes |
| Lab - Golden Cross | M | 1.6 | no |
| Lab - Golden Cross | F | 1.6 | yes |
| Lab - Golden Cross | M | 1.6 | no |
| Lab - Golden Cross | F | 1.6 | yes |
| Lab - Golden Cross | F | 1.6 | yes |
| Lab - Golden Cross | M | 1.6 | no |
| Lab - Golden Cross | M | 1.6 | no |
| Lab - Golden Cross | F | 1.6 | yes |
| Lab - Golden Cross | M | 1.6 | yes |
| Lab - Golden Cross | F | 1.5 | yes |
| Lab - Golden Cross | M | 1.8 | no |
| Labrador Retriever | M | 1.7 | yes |
| Lab - Golden Cross | F | 1.5 | yes |
| Golden Retriever | M | 1.5 | yes |
| Lab - Golden Cross | F | 1.7 | yes |
| Golden Retriever | F | 1.5 | yes |
| Lab - Golden Cross | M | 1.7 | no |
| Labrador Retriever | F | 1.5 | yes |
| Lab - Golden Cross | M | 1.7 | yes |
| Lab - Golden Cross | F | 1.7 | yes |
| Labrador Retriever | M | 1.5 | yes |
| Labrador Retriever | F | 1.5 | yes |
| Labrador Retriever | F | 1.5 | yes |
| Lab - Golden Cross | F | 1.5 | yes |
| Lab - Golden Cross | M | 1.7 | no |
| Lab - Golden Cross | F | 1.7 | yes |
| Lab - Golden Cross | M | 1.6 | no |
| Labrador Retriever | M | 1.4 | yes |

### Procedure

All assistance dogs participated in an initial temperament test (designed and implemented by CCI) which included two social events (threatening stranger and unfamiliar dog) which were video recorded for the purpose of this study. The larger temperament test was administered by walking dogs along a pre-determined path, which included a variety of potentially distracting, startling, or threatening stimuli. Immediately following this temperament test, dogs were also tested with the video stimuli used in Experiment 1. In addition to these behavioral tests, puppy-raisers completed C-BARQ evaluations for the majority of these subjects based on the dog’s behavior at 1 year of age (the C-BARQ could not be obtained for some dogs who were raised in a prison puppy raising program). Blood draws were performed on all dogs as part of a routine veterinary examination one day prior to the behavioral test.

#### Threatening Stranger (TS)

In the threatening stranger test the handler walked the dog toward a man (TS) sitting on a bench who was wearing a hooded jacket and holding a cane. As the dog approached, the TS stood up, banged his cane on the ground, shouted toward the dog in a threatening tone, and walked 2m toward the dog. During the TS’s approach the handler remained stationary with the dog on leash for ~10s to observe the dog’s reaction. The handler then encouraged the dog to approach the man by walking forward while holding the leash. Once the dog was within arm’s reach of the TS, the TS removed his hood, set down his cane, kneeled, and greeted the dog in a friendly manner.

#### Unfamiliar Dog

In the unfamiliar dog trial subjects confronted a three-dimensional lifelike dog model (Old English sheepdog, identical to that from Experiment 1) as they walked (on leash, handled by a trainer) along a sidewalk. This sidewalk wrapped around the exterior of a building such that the model was first became visible at a distance of ~10m. Trainers first led the dogs to a point 6m from the model before turning around and returning to a distance of 10m from the model. Trainers and dogs then made two additional approaches to distances of 3 and 0.6m from the model, each time turning around and walking ~5 steps away from the model before the next approach. On fourth and final approach dogs were allowed to freely inspect the model before advancing to the next item in the temperament test.

#### Video Stimuli

The video stimuli and procedure were identical to Experiment 1.

### Scoring and Analysis

All trials were scored from video by two independent observers.

#### Video Stimuli

Barking, growling, snarling, and lunging were coded as in Experiment 1. However, only barking and growling were observed, and only in a single dog, thus behavioral responses to the video stimuli were excluded from subsequent analysis.

#### Threatening Stranger

From video we coded whether dogs barked, growled, snarled or lunged at the threatening stranger (TS). We also classified the dog’s initial reaction to the TS into one of the following four categories: (1) Dog resisted the handler and was not easily coaxed toward the TS, (2) Dog resisted the handler but was easily coaxed toward the TS, (3) Dog moved toward TS confidently alongside the handler, (4) Dog moved toward TS confidently in front of the handler. Lastly, we used the following classifications for dogs’ reactions to the TS after he removed his hood and greeted the dog (recovery): (1) Dog did not recover from threat and continued to avoid TS after his change in demeanor, (2) Dog remained skittish but was willing to greet and be touched by the TS, (3) Dog was eager to greet TS and showed no signs of fear or hesitation. Inter-rater agreement was excellent for all measures (bark: kappa = 1; growl: kappa = 1; snarl: kappa = 1, initial reaction: R = 0.97, recovery: R = 0.91), but lunging was not observed so was dropped from analysis. For the purpose of analysis, barking, growling and snarling were combined into a single composite aggression score. Because each variable was coded as a binary measure (1: present, 0: absent), the composite aggression score was calculated as the sum of barking, growling and snarling for each dog (range 0-3).

#### Unfamiliar Dog

From video we coded whether dogs barked, growled, or snarled at the unfamiliar dog. We also classified the dog’s approach toward the unfamiliar dog (UD) into one of the following four categories: (1) Dog resisted handler when approaching UD and was not easily coaxed toward UD (2) Dog resisted handler when approaching UD but was easily coaxed toward UD (3) Dog approached UD confidently but was easily redirected away from UD by handler (4) Dog approached UD confidently and was not easily redirected away from UD by handler. There was only one instance of barking and growling, and no instances of snarling, and these variables were therefore dropped prior to analysis. Inter-rater agreement for the approach measure was good (R = 0.88).

#### C-BARQ

As in Experiment 1, we assessed the association between plasma peptide levels and C-BARQ scores for measures related to human- and dog-directed fear and aggression.

#### Statistical Analysis

Due to limited variability and high skew in the behavioral measures, all behavioral measures were discretized into two quantile groups corresponding to low and high scores on each measure using the Hmisc package (Harrell Jr, 2015) in the R Environment for Statistical Computing (R Core Team, 2015). Associations with plasma peptide levels were tested by fitting generalized linear models predicting behavior as a function of OT, AVP and sex. Individual model predictors were evaluated in a model-comparison framework using a likelihood ratio test to determine the change in likelihood when individual variables were added to the model. For population comparisons we included sex as a covariate in the analyses. Hormone data were log transformed prior to analysis.

#### Sample Collection and Hormone Analysis

Samples were collected and processed as described in Experiment 1. Assistance dog samples were initially analyzed for free OT and AVP using the same methods from Experiment 1 to allow direct comparison of free OT and AVP concentrations between assistance and pet dogs. The majority of assistance dog samples (N = 19) were run on the same plates as pet dog samples allowing a direct comparison between these groups. Eleven assistance dog samples were run on a plate containing no pet dog samples, so to control for inter-assay variance we excluded these samples from the population comparison for free OT and AVP. For comparison within the assistance dog group, all samples were re-run on the same plate to allow the most direct comparisons within this population. For comparisons of free OT and AVP within the assistance dog population, we adopted a modified extraction protocol which yielded improved OT and AVP recovery (SOM). OT samples for this analysis were also measured using a different ELISA kit, which permitted detection in a better region of the kit’s standard curve (see SOM for validation). For comparison of total OT and AVP all pet and assistance dog samples were initially run on the same plates allowing all individuals to be included in the population comparison. As above, all assistance dog samples were then rerun on the same plate allowing the most direct comparisons within this population.

## Results

### Population Comparison

Comparison of pet and candidate assistance dog samples revealed a population difference for both free and total OT (free OT: χ 2 =4.50, df = 1, p = 0.03; total OT: χ 2 =16.93, df = 1, p < 0.001) but not for either measure of AVP (free AVP: χ 2 = 2.01, df = 1, p = 0.16, total AVP: χ 2 = 1.53, df = 1, p = 0.22). Specifically, the candidate assistance dogs – who were systematically bred for calm temperaments and friendly demeanors - had higher free and total plasma OT than pet dogs (Figure 4). Additionally, assistance dogs had higher free and total OT:AVP ratios then pet dogs (Figure 4; free OT:AVP: χ 2 = 4.01, df = 1, p = 0.05, total OT:AVP: χ 2 = 5.33, df = 1, p = 0.02).

**Figure 4.**
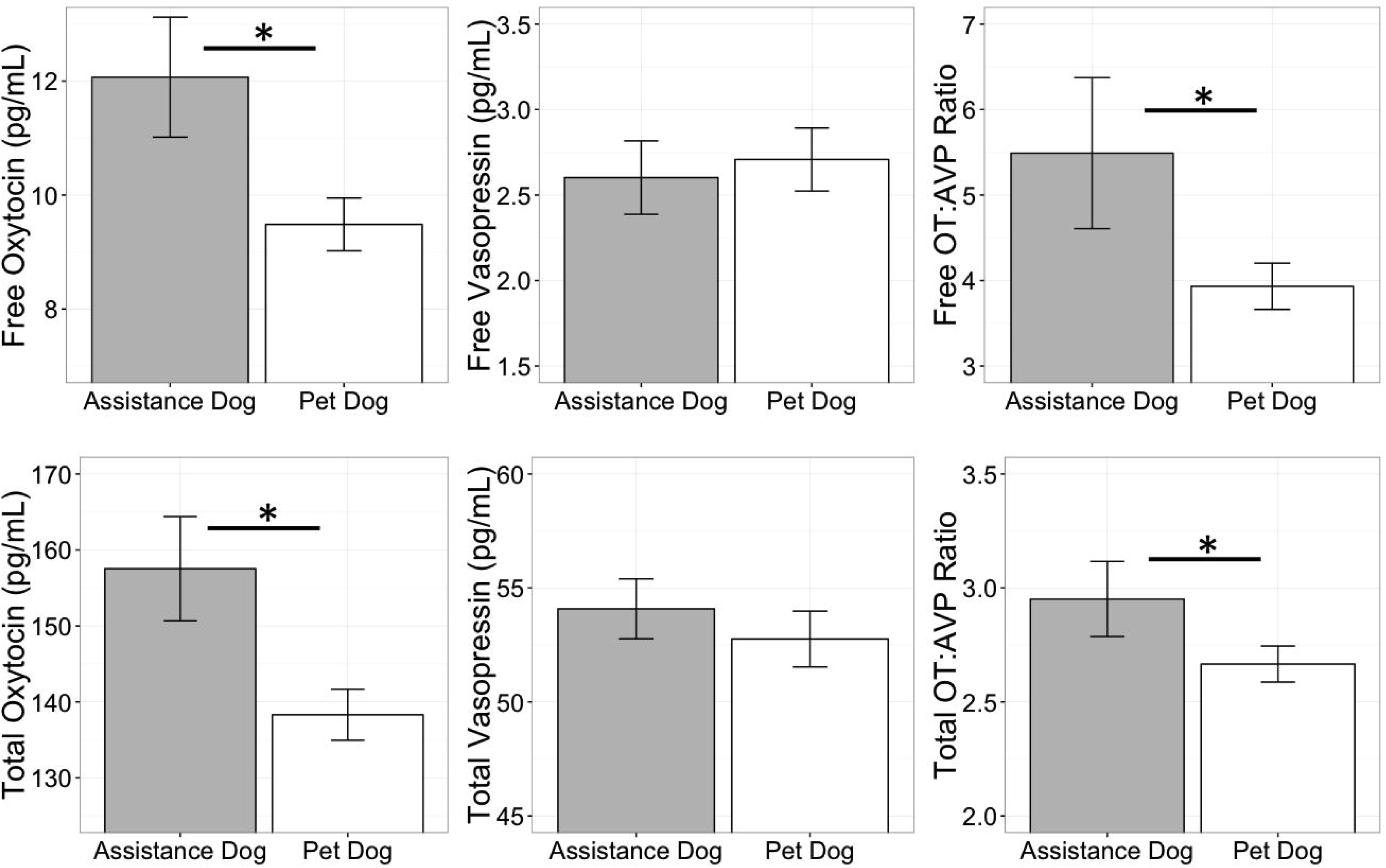

### Threatening Stranger and Unfamiliar Dog

Hormonal associations with behavior in the unfamiliar dog and threatening stranger test are summarized in Table 4. Although dogs varied in their willingness to approach the unfamiliar dog and threatening stranger, there were few cases of aggressive behaviors. In the threatening stranger test, only a minority of subjects (~20%) barked, growled, or snarled at the stranger, and there were no significant associations between these behaviors and free OT, AVP, or the OT:AVP ratio (Table 4). There were no associations between the free hormone measures and initial reactions to the threatening stranger (Table 4), however dogs who remained skittish or were hesitant to greet the stranger after he changed demeanor had higher free OT:AVP ratios (low AVP relative to OT). Analysis of total peptide concentrations revealed that dogs who behaved aggressively toward the threatening stranger had significantly higher total AVP than dogs who did not, mirroring the patterns observed in Experiment 1 (Table 4). Dogs who behaved aggressively in this context also exhibited significantly lower OT:AVP ratios than dogs who did not (Table 4). Lastly, in the unfamiliar dog test, dogs who exhibited the most confident approaches toward the model dog were characterized by higher free OT:AVP ratios than dogs who were more reluctant to approach, and there were no other significant associations (Table 4).

**Table 4.** Associations between free and total oxytocin and vasopressin and behavior during the unfamiliar dog and threatening stranger test. Significant associations are bolded.

|  | Free Peptide Concentration |  |  |  |  |  | Total Peptide Concentration |  |  |  |  |  |
| --- | --- | --- | --- | --- | --- | --- | --- | --- | --- | --- | --- | --- |
|  | AVP |  | OT |  | OT:AVP Ratio |  | AVP |  | OT |  | OT:AVP Ratio |  |
| | $\chi^2$ | p | $\chi^2$ | p | $\chi^2$ | p | $\chi^2$ | p | $\chi^2$ | p | $\chi^2$ | p |
| Unfamiliar Dog: Approach | 1.25 | 0.26 | 2.49 | 0.11 | 4.56 | **0.03** | 2.73 | 0.1 | 0.81 | 0.37 | 0.01 | 0.91 |
| Threatening Stranger: Aggression | 2.22 | 0.14 | 0.15 | 0.70 | 1.83 | 0.18 | 6.74 | **0.01** | 0.55 | 0.46 | 4.38 | **0.04** |
| Threatening Stranger: Initial Reaction | 1.41 | 0.23 | 0.84 | 0.36 | 1.32 | 0.25 | 0.38 | 0.54 | 1.62 | 0.2 | 2.05 | 0.15 |
| Threatening Stranger: Recovery | 2.30 | 0.13 | 1.32 | 0.25 | 3.89 | **0.05** | 0.67 | 0.41 | 2.11 | 0.15 | 3.22 | 0.07 |

#### C-BARQ

Within the assistance dog population there was minimal variation on all C-BARQ measures with the exception of dog-directed fear, which was negatively associated with the free OT:AVP ratio (SOM).

## Discussion

The results of Experiment 2 are consistent with, and build on the findings from Experiment 1. Although there was minimal aggressive behavior in this sample, dogs who behaved aggressively toward the threatening stranger had higher total AVP concentrations, and higher total AVP relative to OT, corroborating the patterns observed in Experiment 1. We also observed a population difference in OT, but not AVP concentrations between the pet dog population from Experiment 1, and the population of candidate assistance dogs. Specifically, candidate assistance dogs had higher free and total plasma OT than pet dogs, as well as a higher ratio of OT to AVP on both measures. Given that this population has been actively selected for calm, affiliative and non-aggressive temperaments for more than 40 years, one possibility is that this phenotype has been achieved in part due to upregulation of the oxytocinergic system. This possibility is supported by data from dog OT administration studies, in which exogenous OT has been shown to increase affiliative behavior (Nagasawa et al., 2015; Romero et al., 2014; but see Hernádi et al. 2015), and promote calm emotional states by decreasing heart rate and increasing heart-rate variability (Kovács et al., 2016). However, this population difference should be interpreted cautiously as several factors varied between the assistance and pet dog populations. Specifically, whereas the pet dog population was heterogeneous with respect to breed, age and body size, the assistance dog population consisted entirely of young retriever dogs. Given potential for breed differences in OT genetics (Bence et al., 2016) future investigations of possible differences between pet and selectively bred working populations will benefit from designs controlling for these parameters.

## General Discussion

Aggression is a complex and multifaceted construct that is likely to have an equally complex etiology (Blanchard, Wall, & Blanchard, 2003). Aggressive behaviors may be influenced by a wide range of genetic, epigenetic, and environmental factors, as well as an interrelated network of hormonal and physiological processes. Here we report the first systematic investigation of the relationships between endogenous oxytocin (OT), vasopressin (AVP), and aggression in domestic dogs. Our main findings were associations between plasma AVP concentrations and aggressive behavior toward both conspecifics and humans, but the direction of these relationships varied depending on whether we considered only the free fraction, or total AVP (free and bound). In Experiment 1, pet dogs with a history of chronic aggression toward conspecifics had lower free AVP, but higher total AVP than a group of controls matched on breed, sex, and age. In Experiment 2, candidate assistance dogs who behaved more aggressively toward a threatening (human) stranger had higher total AVP than dogs who exhibited less aggression in this context. Therefore, across studies we observed a consistent positive association between total AVP concentrations and aggressive behavior, with some evidence for an opposite association with respect to free AVP.

The difference in results between these two measurement approaches adds a layer of complexity to ongoing debates regarding the most appropriate methods for quantification of OT and AVP in plasma (Brandtzaeg et al., 2016; Leng & Sabatier, 2016; Martin & Carter, 2013; Szeto et al., 2011). Historically, researchers have measured these peptides either in (non-extracted) diluted plasma, or following solid phase extraction (SPE). The former approach tends to yield much higher concentrations, but may suffer from matrix interference, as we observed in our validation studies. The latter approach reduces these sources of interference, but yields concentrations near the lower limit of detection, in part due to poor recovery of the target analyte (Brandtzaeg et al., 2016). However, it has recently been shown that OT (and likely AVP) binds strongly to plasma proteins, and that breaking these bonds (followed by protein precipitation) allows for quantification of total OT and AVP, which are present in plasma at remarkably high concentrations (Brandtzaeg et al., 2016). Therefore, this approach eliminates the most common problems associated with SPE (concentrations near the lower limit of detection) and the analysis of non-extracted plasma (interference from plasma proteins).

As biomarkers, the significance of free vs. total peptide concentrations remains poorly understood. One possibility is that total OT/AVP may provide the most powerful approach for comparing basal individual differences, indexing long-term peptidergic activity. In contrast, free OT/AVP may be a more useful metric for assessing acute responses to stimuli. Therefore, higher total plasma AVP in dogs prone to aggression may reflect general overactivity in the vasopressinergic system, despite the fact that these dogs had lower levels of free AVP before the behavioral test. This possibility is consistent with data implicating AVP as a primary activator of the hypothalamic-pituitary-adrenal (HPA) axis (Aguilera & Rabadan-Diehl, 2000; Scott & Dinan, 1998), which has long been known to facilitate ‘fight or flight’ behavior (Cannon, 1932; Kruk, Halasz, Meelis, & Haller, 2004). The positive association between AVP and aggression is also consistent with previous studies in humans, which have revealed a positive correlation between AVP in cerebrospinal fluid and a life history of aggression, as well as animal studies documenting that centrally administered AVP facilitates aggression toward conspecifics, whereas AVP antagonists diminish this response (Ferris, 1992; Ferris et al., 2006).

However, a wealth of data suggests that the functions of AVP and OT may be highly dependent on species and sex-specific factors, as well as the site of action in the brain. For example, studies that have measured local AVP release, or manipulated the AVP system through receptor agonism/antagonism reveal that AVP can both facilitate and inhibit aggression (De Boer, Olivier, Veening, & Koolhaas, 2015; Kelly & Goodson, 2014; Veenema, Beiderbeck, Lukas, & Neumann, 2010). Similarly, AVP concentrations and the density of AVP tracts in various regions of the brain have also been associated with aggression, but the nature of these relationships has been variable between species. In mice selected for short or long attack latencies, and high and low aggression wild-type rats, less aggressive individuals are characterized by greater vasopressinergic immunoreactivity in several brain regions (Compaan, Buijs, Pool, & De Ruiter, 1993; Everts, De Ruiter, & Koolhaas, 1997). However, contrary results have been reported in comparisons of other rodent species suggesting that relationships between AVP and aggression are unlikely to be uniform across mammalian taxa (De Boer et al., 2015).

Studies of peripheral peptide concentrations are further complicated by a lack of knowledge regarding the correspondence between central and peripheral peptide release. Although some studies have reported positive correlations between OT and AVP concentrations in cerebrospinal fluid (CSF) and free OT/AVP in plasma, others have failed to detect this relationship (Carson et al., 2015; Carson et al., 2014; Kagerbauer et al., 2013). Studies of peptide release patterns have been similarly mixed with evidence for both correlated, and independent central and peripheral release (Neumann & Landgraf, 2012). Lastly, the ultimate effects of these peptides depend not only on their concentrations, but also in the distribution and state of their receptors, which vary not only between individuals and species, but also within individuals as a result of other hormonal and epigenetic processes (Carter & Porges, 2013).

Although we found no direct associations between OT and aggression, the population of assistance dogs – which has been systematically selected for affiliative and non-aggressive temperaments – had higher free and total OT than the pet dog population. Given that OT has been positively associated with affiliative social behaviors, these findings are consistent with the possibility that the behavioral targets of selection in the assistance dog population are supported by OT-related mechanisms. However, the assistance and pet dog populations were not matched on critical parameters such as age, breed, or sterilization status, and future research will benefit by controlling for these factors.

At the behavioral level, these studies reveal that realistic three-dimensional models are a useful tool for evaluating dog aggression. Dogs recruited for a history of aggression toward other dogs were much more likely to behave aggressively toward these models than matched controls, and across studies, the three-dimensional models elicited stronger reactions than video-projected stimuli. Importantly, responses to these models were not generalized reactions to novel objects, as dogs reacted much less frequently to comparably-sized control stimuli. Therefore, while unlikely to provide a perfect proxy for real-world behavior (Shabelansky et al., 2015), working with life-like models provides a highly-feasible, safe, and ethical option for assessing aggression, and may also be useful for systematic desensitization with dogs prone to aggressive behavior. In contrast, although several studies have successfully used video-projected stimuli with dogs (e.g., Pongrácz, Miklósi, Dóka, & Csányi, 2003), our video stimuli uniformly elicited minimal response. Importantly, these video stimuli were projected to life-like dimensions, and presented with a refresh rate suitable for the flicker-fusion frequency of dog vision (Coile, Pollitz, & Smith, 1989). Thus while dogs were likely capable of perceiving the video, three-dimensional cues may be required for successfully simulating an encounter with a conspecific.

Collectively, our findings provide preliminary evidence that AVP may be an important mediator of canine aggression and call for further research in this area. Because previous studies have focused primarily on the role of serotonin and testosterone in dog aggression – both of which contribute to pathways that also include OT and AVP (Delville, Mansour, & Ferris, 1996; Dölen, Darvishzadeh, Huang, & Malenka, 2013) – future research will benefit by addressing the complex interactions between these systems (Weisman & Feldman, 2013). Similarly, several exploratory studies have begun to investigate the effects of intranasal peptide administration in dogs, which in some cases has led to notable changes in social behavior and cognition (Hernádi et al., 2015; Oliva et al., 2015; Oliva, Rault, Appleton, & Lill, 2016; Romero et al., 2014). Therefore, OT/AVP administration provides another non-invasive possibility for investigating how OT and AVP may modulate aggressive behavior in dogs. Notably, previous studies using this approach have found no evidence that intranasal OT decreases aggressive behavior in dogs (Hernádi et al., 2015), which is consistent with our findings that plasma OT levels were unrelated to individual differences in aggression.

Given that we found evidence for both positive and negative associations between AVP and aggression, we have no strong predictions regarding how AVP administration may influence aggressive behavior. Historically, AVP has been noted to have effects antagonistic to those of OT, increasing anxiety, depression and stress responses (Benarroch, 2013). However, several studies reveal that AVP administration can have effects very similar to those of OT, decreasing heart rate (Hicks et al., 2014), reducing anxiety (Appenrodt, Schnabel, & Schwarzberg, 1998) and increasing sociability (Ramos, Hicks, Caminer, & McGregor, 2014). Notably, many of the prosocial effects of OT, AVP and MDMA (‘Ecstasy’) are believed to act through AVP networks in the brain (Ramos et al., 2013), and blocking AVP, but not OT receptors inhibits many of the effects of both OT and AVP, suggesting that the actions of both peptides are dependent on an AVP receptor-dependent mechanism (Hicks et al., 2014). Therefore, as with OT, the effects of AVP administration are likely to be complex, and moderated by a diverse range of biological factors (Bartz, Zaki, Bolger, & Ochsner, 2011).

Ultimately, dog aggression is a normal and adaptive social behavior, but expressed in the wrong contexts, or to an extreme extent, its consequences jeopardize the welfare of both humans and dogs in our society. Our research reveals potential links between the neuropeptides oxytocin and vasopressin and aggression in dogs, and sets the stage for future work to evaluate whether treatments and interventions for aggression can be improved by considering the roles of these peptides. Ultimately we hope that these investigations will lead to increased knowledge of the biology of aggression, promote human and animal welfare, and help to preserve the unique and long-standing relationship between humans and dogs.

## Acknowledgements

We thank Eliot Cohen, Kerri Rodriquez, Lyndy Bleu Harden Plumley, Alexa King, Janet Bogan, Kerinne Levy, and Brenda Kennedy for help with data collection. We thank Duncan Lascelles and the Clinical Services Core at North Carolina State University College of Veterinary Medicine for their support of this research. We thank James Serpell and the University of Pennsylvania for allowing us to use the C-BARQ as a part of this study, and DOGTV for allowing us to use video footage for the purpose of these experiments. We gratefully acknowledge the Stanton Foundation for a Next Generation Canine Research Fellowship to ELM, which supported this research. Grants from the Fetzer Foundation and NIH (P01 HD 075750; C.S. Carter, PI) sponsored much of the preliminary work on the measurement of oxytocin which provided background for this research.

## References

Aguilera, G., & Rabadan-Diehl, C. (2000). Vasopressinergic regulation of the hypothalamic– pituitary–adrenal axis: implications for stress adaptation. Regulatory peptides, 96(1), 23–29.

Albers, H. E. (2012). The regulation of social recognition, social communication and aggression: vasopressin in the social behavior neural network. Hormones and Behavior, 61(3), 283–292.

Albers, H. E. (2015). Species, sex and individual differences in the vasotocin/vasopressin system: relationship to neurochemical signaling in the social behavior neural network. Frontiers in neuroendocrinology, 36, 49–71.

Amat, M., Le Brech, S., Camps, T., Torrente, C., Mariotti, V. M., Ruiz, J. L., et al. (2013). Differences in serotonin serum concentration between aggressive English cocker spaniels and aggressive dogs of other breeds. Journal of Veterinary Behavior: Clinical Applications and Research, 8(1), 19–25.

Appenrodt, E., Schnabel, R., & Schwarzberg, H. (1998). Vasopressin administration modulates anxiety-related behavior in rats. Physiology & behavior, 64(4), 543–547.

Archer, J. (1988). The behavioural biology of aggression (Vol. 1): CUP Archive.

Bartz, J. A., Zaki, J., Bolger, N., & Ochsner, K. N. (2011). Social effects of oxytocin in humans: context and person matter. Trends in cognitive sciences, 15(7), 301–309.

Beetz, A., Uvnäs-Moberg, K., Julius, H., & Kotrschal, K. (2012). Psychosocial and psychophysiological effects of human-animal interactions: the possible role of oxytocin. Frontiers in Psychology, 3.

Benarroch, E. E. (2013). Oxytocin and vasopressin Social neuropeptides with complex neuromodulatory functions. Neurology, 80(16), 1521–1528.

Bence, M., Marx, P., Szantai, E., Kubinyi, E., Ronai, Z., & Banlaki, Z. (2016). Lessons from the canine Oxtr gene: populations, variants and functional aspects. Genes, Brain and Behavior.

Bester - Meredith, J. K., Martin, P. A., & Marler, C. A. (2005). Manipulations of vasopressin alter aggression differently across testing conditions in monogamous and non - monogamous Peromyscus mice. Aggressive Behavior, 31(2),189–199.

Blanchard, R. J., Wall, P. M., & Blanchard, D. C. (2003). Problems in the study of rodent aggression. Hormones and behavior, 44(3), 161–170.

Brandtzaeg, O. K., Johnsen, E., Roberg-Larsen, H., Seip, K. F., MacLean, E. L., Gesquiere, L. R., et al. (2016). Proteomics tools reveal startlingly high amounts of oxytocin in plasma and serum. Scientific Reports.

Caldwell, H. K., Lee, H.-J., Macbeth, A. H., & Young, III W. S. (2008). Vasopressin: behavioral roles of an “original” neuropeptide. Progress in Neurobiology, 84(1), 1–24.

Cannon, W. B. (1932). The wisdom of the body.

Carson, D., Berquist, S., Trujillo, T., Garner, J., Hannah, S., Hyde, S., et al. (2015). Cerebrospinal fluid and plasma oxytocin concentrations are positively correlated and negatively predict anxiety in children. Molecular psychiatry, 20(9), 1085–1090.

Carson, D. S., Howerton, C. L., Garner, J. P., Hyde, S. A., Clark, C. L., Hardan, A. Y., et al. (2014). Plasma vasopressin concentrations positively predict cerebrospinal fluid vasopressin concentrations in human neonates. Peptides, 61, 12–16.

Carter, C. S. (1998). Neuroendocrine perspectives on social attachment and love. Psychoneuroendocrinology, 23(8), 779–818.

Carter, C. S. (2014). Oxytocin pathways and the evolution of human behavior. Annual review of psychology, 65, 17–39.

Carter, C. S., Devries, A. C., & Getz, L. L. (1995). Physiological substrates of mammalian monogamy: the prairie vole model. Neuroscience & Biobehavioral Reviews, 19(2), 303–314.

Carter, C. S., Grippo, A. J., Pournajafi-Nazarloo, H., Ruscio, M. G., & Porges, S. W. (2008). Oxytocin, vasopressin and sociality. In D. N. Inga & L. Rainer (Eds.), Progress in Brain Research (Vol. Volume 170, pp. 331–336): Elsevier.

Carter, C. S., & Porges, S. W. (2013). The biochemistry of love: an oxytocin hypothesis. EMBO reports, 14(1), 12–16.

Centers for Disease Control and Prevention. (2003). Nonfatal dog bite-related injuries treated in hospital emergency departments–United States, 2001. MMWR: Morbidity and Mortality Weekly Report, 52(26), 605–610.

Christensen, J. C., Shiyanov, P. A., Estepp, J. R., & Schlager, J. J. (2014). Lack of association between human plasma oxytocin and interpersonal trust in a prisoner’s dilemma paradigm. PLoS one, 9(12), e116172.

Coccaro, E. F., Kavoussi, R. J., Hauger, R. L., Cooper, T. B., & Ferris, C. F. (1998). Cerebrospinal fluid vasopressin levels correlates with aggression and serotonin function in personality-disordered subjects. Archives of General Psychiatry, 55(8), 708–714.

Coile, D. C., Pollitz, C. H., & Smith, J. C. (1989). Behavioral determination of critical flicker fusion in dogs. Physiol Behav, 45(6), 1087–1092.

Compaan, J., Buijs, R., Pool, C., & De Ruiter, A. (1993). Differential lateral septal vasopressin innervation in aggressive and nonaggressive male mice. Brain research bulletin, 30(1), 1–6.

De Boer, S., Olivier, B., Veening, J., & Koolhaas, J. (2015). The neurobiology of offensive aggression: Revealing a modular view. Physiology & behavior, 146, 111–127.

Delville, Y., Mansour, K. M., & Ferris, C. F. (1996). Testosterone facilitates aggression by modulating vasopressin receptors in the hypothalamus. Physiology & Behavior, 60(1), 25–29.

Dölen, G., Darvishzadeh, A., Huang, K. W., & Malenka, R. C. (2013). Social reward requires coordinated activity of nucleus accumbens oxytocin and serotonin. Nature, 501(7466), 179–184.

Donaldson, Z. R., & Young, L. J. (2008). Oxytocin, vasopressin, and the neurogenetics of sociality. Science, 322(5903), 900–904.

Duffy, D. L., Hsu, Y., & Serpell, J. A. (2008). Breed differences in canine aggression. Applied Animal Behaviour Science, 114(3), 441–460.

Everts, H. G., De Ruiter, A., & Koolhaas, J. M. (1997). Differential lateral septal vasopressin in wild-type rats: correlation with aggression. Hormones and behavior, 31(2), 136–144.

Ferris, C. (1992). Role of vasopressin in aggressive and dominant/subordinate behaviors. Annals of the New York Academy of Sciences, 652(1), 212–226.

Ferris, C. F., Lu, S.-f., Messenger, T., Guillon, C. D., Heindel, N., Miller, M., et al. (2006). Orally active vasopressin V1a receptor antagonist, SRX251, selectively blocks aggressive behavior. Pharmacology Biochemistry and Behavior, 83(2), 169–174.

Ferris, C. F., Melloni Jr, R. H., Koppel, G., Perry, K. W., Fuller, R. W., & Delville, Y. (1997). Vasopressin/serotonin interactions in the anterior hypothalamus control aggressive behavior in golden hamsters. The Journal of Neuroscience, 17(11), 4331–4340.

Ferris, C. F., & Potegal, M. (1988). Vasopressin receptor blockade in the anterior hypothalamus suppresses aggression in hamsters. Physiology & behavior, 44(2), 235–239.

Gail, M. H., Lubin, J. H., & Rubinstein, L. V. (1981). Likelihood calculations for matched casecontrol studies and survival studies with tied death times. Biometrika, 68(3), 703–707.

Gilchrist, J., Sacks, J., White, D., & Kresnow, M. (2008). Dog bites: still a problem? Injury Prevention, 14(5), 296–301.

Gobrogge, K. L., Liu, Y., Jia, X., & Wang, Z. (2007). Anterior hypothalamic neural activation and neurochemical associations with aggression in pair - bonded male prairie voles. Journal of Comparative Neurology, 502(6),1109–1122.

Goodson, J. L., & Bass, A. H. (2001). Social behavior functions and related anatomical characteristics of vasotocin/vasopressin systems in vertebrates. Brain Research Reviews, 35(3), 246–265.

Guy, N. C., Luescher, U., Dohoo, S. E., Spangler, E., Miller, J. B., Dohoo, I. R., et al. (2001). Demographic and aggressive characteristics of dogs in a general veterinary caseload. Applied Animal Behaviour Science, 74(1), 15–28.

Harrell Jr, F. E. (2015). Hmisc: Harrell Miscellaneous. R package version 3.17–1.

Haug, L. I. (2008). Canine aggression toward unfamiliar people and dogs. Veterinary Clinics of North America: Small Animal Practice, 38(5),1023–1041.

Hernádi, A., Kis, A., Kanizsár, O., Tóth, K., Miklósi, B., & Topál, J. (2015). Intranasally administered oxytocin affects how dogs (Canis familiaris) react to the threatening approach of their owner and an unfamiliar experimenter. Behavioural processes, 119, 1–5.

Hicks, C., Ramos, L., Reekie, T., Misagh, G., Narlawar, R., Kassiou, M., et al. (2014). Body temperature and cardiac changes induced by peripherally administered oxytocin, vasopressin and the non - peptide oxytocin receptor agonist WAY 267,464: a biotelemetry study in rats. British journal of pharmacology, 171(11), 2868–2887.

Johnsen, E., Leknes, S., Wilson, S. R., & Lundanes, E. (2015). Liquid chromatography-mass spectrometry platform for both small neurotransmitters and neuropeptides in blood, with automatic and robust solid phase extraction. Scientific reports, 5.

Kagerbauer, S., Martin, J., Schuster, T., Blobner, M., Kochs, E., & Landgraf, R. (2013). Plasma oxytocin and vasopressin do not predict neuropeptide concentrations in human cerebrospinal fluid. Journal of neuroendocrinology, 25(7), 668–673.

Kelly, A. M., & Goodson, J. L. (2014). Social functions of individual vasopressin–oxytocin cell groups in vertebrates: What do we really know? Frontiers in neuroendocrinology.

Kis, A., Bence, M., Lakatos, G., Pergel, E., Turcsán, B., Pluijmakers, J., et al. (2014). Oxytocin Receptor Gene Polymorphisms Are Associated with Human Directed Social Behavior in Dogs (*Canis familiaris*). PloS one, 9(1), e83993.

Kovács, K., Kis, A., Kanizsár, O., Hernádi, A., Gácsi, M., & Topál, J. (2016). The effect of oxytocin on biological motion perception in dogs (Canis familiaris). Animal cognition, 19(3), 513–522.

Kruk, M. R., Halasz, J., Meelis, W., & Haller, J. (2004). Fast positive feedback between the adrenocortical stress response and a brain mechanism involved in aggressive behavior. Behavioral neuroscience, 118(5), 1062.

Leng, G., & Sabatier, N. (2016). Measuring oxytocin and vasopressin: bioassays, immunoassays and random numbers. Journal of Neuroendocrinology.

León, M., Rosado, B., García-Belenguer, S., Chacón, G., Villegas, A., & Palacio, J. (2012). Assessment of serotonin in serum, plasma, and platelets of aggressive dogs. Journal of Veterinary Behavior: Clinical Applications and Research, 7(6), 348–352.

MacLean, E. L., & Hare, B. (2015). Dogs hijack the human bonding pathway. Science, 348(6232), 280–281.

MacLean, E. L., Herrmann, E., Suchindran, S., & Hare, B. (2017). Individual differences in cooperative communicative skills are more similar between dogs and humans than chimpanzees. Animal Behaviour, 126, 41–51.

Martin, W., & Carter, C. S. (2013). Oxytocin and vasopressin are sequestered in plasma. Paper presented at the World Congress of Neurohypophyseal Hormones Abstracts. Bristol, England.

Martin, W. L. (2014). Int. Patent Pub No: WO/2014/210399.

Mitsui, S., Yamamoto, M., Nagasawa, M., Mogi, K., Kikusui, T., Ohtani, N., et al. (2011). Urinary oxytocin as a noninvasive biomarker of positive emotion in dogs. Hormones and behavior, 60(3), 239–243.

Nagasawa, M., Mitsui, S., En, S., Ohtani, N., Ohta, M., Sakuma, Y., et al. (2015). Oxytocin-gaze positive loop and the coevolution of human-dog bonds. Science, 348(6232), 333–336.

Neilson, J. C., Eckstein, R. A., & Hart, B. (1997). Effects of castration on problem behaviors in male dogs with reference to age and duration of behavior. Journal of the American Veterinary Medical Association, 211(2), 180–182.

Neumann, I. D., & Landgraf, R. (2012). Balance of brain oxytocin and vasopressin: implications for anxiety, depression, and social behaviors. Trends in neurosciences, 35(11), 649–659.

Odendaal, J., & Meintjes, R. (2003). Neurophysiological correlates of affiliative behaviour between humans and dogs. The Veterinary Journal, 165(3), 296–301.

Oliva, J., Rault, J.-L., Appleton, B., & Lill, A. (2015). Oxytocin enhances the appropriate use of human social cues by the domestic dog (*Canis familiaris*) in an object choice task. Animal cognition, 1–9.

Oliva, J. L., Rault, J.-L., Appleton, B., & Lill, A. (2016). Oxytocin blocks pet dog (Canis familiaris) object choice task performance being predicted by owner-perceived intelligence and owner attachment. Pet Behaviour Science(1), 31–46.

Oliva, J. L., Wong, Y. T., Rault, J.-L., Appleton, B., & Lill, A. (2016). The oxytocin receptor gene, an integral piece of the evolution of Canis familaris from Canis lupus. Pet Behaviour Science(2), 1–15.

Patronek, G. J., Glickman, L. T., Beck, A. M., McCabe, G. P., & Ecker, C. (1996). Risk factors for relinquishment of dogs to an animal shelter. Journal of the American Veterinary Medical Association, 209(3), 572–581.

Pongrácz, P., Miklósi, Á., Dóka, A., & Csányi, V. (2003). Successful Application of Video **-** Projected Human Images for Signalling to Dogs. Ethology, 109(10), 809–821.

R Core Team. (2015). R: A Language and Environment for Statistical Computing. Vienna, Austria: R Foundation for Statistical Computing.

Ramos, L., Hicks, C., Caminer, A., & McGregor, I. S. (2014). Inhaled vasopressin increases sociability and reduces body temperature and heart rate in rats. Psychoneuroendocrinology, 46, 46–51.

Ramos, L., Hicks, C., Kevin, R., Caminer, A., Narlawar, R., Kassiou, M., et al. (2013). Acute prosocial effects of oxytocin and vasopressin when given alone or in combination with 3, 4-methylenedioxymethamphetamine in rats: involvement of the V1A receptor. Neuropsychopharmacology, 38(11), 2249–2259.

Rehn, T., Handlin, L., Uvnäs-Moberg, K., & Keeling, L. J. (2014). Dogs’ endocrine and behavioural responses at reunion are affected by how the human initiates contact. Physiology & Behavior, 124, 45–53.

Reisner, I. R., Mann, J. J., Stanley, M., Huang, Y., & Houpt, K. A. (1996). Comparison of cerebrospinal fluid monoamine metabolite levels in dominant-aggressive and non-aggressive dogs. Brain research, 714(1), 57–64.

Robinson, K. J., Hazon, N., Lonergan, M., & Pomeroy, P. P. (2014). Validation of an enzyme-linked immunoassay (ELISA) for plasma oxytocin in a novel mammal species reveals potential errors induced by sampling procedure. Journal of neuroscience methods, 226, 73–79.

Romero, T., Nagasawa, M., Mogi, K., Hasegawa, T., & Kikusui, T. (2014). Oxytocin promotes social bonding in dogs. Proceedings of the National Academy of Sciences, 201322868.

Rosado, B., García-Belenguer, S., León, M., Chacón, G., Villegas, A., & Palacio, J. (2010). Blood concentrations of serotonin, cortisol and dehydroepiandrosterone in aggressive dogs. Applied Animal Behaviour Science, 123(3), 124–130.

Salman, M., New, J., John G, Scarlett, J. M., Kass, P. H., Ruch-Gallie, R., & Hetts, S. (1998). Human and animal factors related to relinquishment of dogs and cats in 12 selected animal shelters in the United States. Journal of Applied Animal Welfare Science, 1(3), 207–226.

Scott, L. V., & Dinan, T. G. (1998). Vasopressin and the regulation of hypothalamic-pituitary-adrenal axis function: implications for the pathophysiology of depression. Life sciences, 62(22), 1985–1998.

Shabelansky, A., Dowling-Guyer, S., Quist, H., D’Arpino, S. S., & McCobb, E. (2015). Consistency of shelter dogs’ behavior toward a fake versus real stimulus dog during a behavior evaluation. Applied Animal Behaviour Science, 163, 158–166.

Szeto, A., McCabe, P. M., Nation, D. A., Tabak, B. A., Rossetti, M. A., McCullough, M. E., et al. (2011). Evaluation of enzyme immunoassay and radioimmunoassay methods for the measurement of plasma oxytocin. Psychosomatic Medicine, 73(5), 393.

Thompson, R., George, K., Walton, J., Orr, S., & Benson, J. (2006). Sex-specific influences of vasopressin on human social communication. Proceedings of the National Academy of Sciences, 103(20), 7889–7894.

van den Berg, S. M., Heuven, H., van den Berg, L., Duffy, D. L., & Serpell, J. A. (2010). Evaluation of the C-BARQ as a measure of stranger-directed aggression in three common dog breeds. Applied animal behaviour science, 124(3), 136–141.

Veenema, A. H., Beiderbeck, D. I., Lukas, M., & Neumann, I. D. (2010). Distinct correlations of vasopressin release within the lateral septum and the bed nucleus of the stria terminalis with the display of intermale aggression. Hormones and behavior, 58(2), 273–281.

Weisman, O., & Feldman, R. (2013). Oxytocin administration affects the production of multiple hormones. Psychoneuroendocrinology, 38(5), 626.

Zhang, G., Zhang, Y., Fast, D. M., Lin, Z., & Steenwyk, R. (2011). Ultra sensitive quantitation of endogenous oxytocin in rat and human plasma using a two-dimensional liquid chromatography–tandem mass spectrometry assay. Analytical biochemistry, 416(1), 45–52.

